# Decentralized Mixed Effects Modeling in COINSTAC

**DOI:** 10.1101/2023.05.12.540598

**Authors:** Sunitha Basodi, Rajikha Raja, Harshvardhan Gazula, Javier Tomas Romero, Sandeep Panta, Thomas Maullin-Sapey, Thomas Nichols, Vince D. Calhoun

## Abstract

Performing group analysis on magnetic resonance imaging (MRI) data with linear mixed-effects (LME) models is challenging due to its large dimensionality and inherent multi-level covariance structure. In addition, as large-scale collaborative projects become commonplace in neuroimaging, data must increasingly be stored and analysed at different locations. In such settings, substantial overheads occur in terms of data transfer and coordination between participating research groups. In some cases, data cannot be pooled together due to privacy or regulatory concerns. In this work, we propose a decentralized LME model to perform a large-scale analysis of data from different collaborations without sharing or pooling. This method is efficient as it overcomes the hurdles of data privacy for sharing and has lower bandwidth and memory requirements for analysis than the centralized modeling approach. We evaluate our model using features extracted from structural magnetic resonance imaging (*s*MRI) data. Results highlight gray matter reductions in the temporal lobe/insula and medical front regions demonstrate the correctness of decentralized LME models. Our analysis also demonstrates that decentralized LME models achieve similar performance compared to the models trained with all the data in one location. We also implement the decentralized LME approach in COINSTAC, a decentralized platform for federating neuroimaging analysis, to demonstrate its value to the neuroimaging community.

## 1 Introduction

Linear mixed-effects (LME) models are commonly used in human neuroscience research to determine the relationship between predictors and outcomes, while accounting for grouping factors and covariance structure present in the data [1, 2, 3, 4, 5, 6, 7, 8, 9]. Modelling LME is computationally challenging in large dimensional brain data especially dealing with structural magnetic resonance imaging (sMRI) or functional MRI (fMRI) data as they typically involve running not one LME, but rather tens of thousands, each corresponding to a voxel in an image (or a freesurfer feature). Further such analyses are theoretically complex, as grouping structure present in the data requires careful attention and iterative computation. Several studies have used this approach to perform group analysis on fMRI data and determine underlying feature relationships, variance-covariance structures of random effects [10], [11], [12], [13]. Though several studies are performing such analyses, these tend to be smaller-scale datasets, typically ranging from 10 to 100 subjects [14], due to the expensive data collection process and lack of availability of many subjects with MRI data, as one is limited by the amount of data that can be gathered for analysis.

To combat the reduced power that arises from small sample sizes, large-scale population datasets, such as the Adolescent Brain Cognitive Development (ABCD) [15] and the UK Biobank study[16] which include imaging data of thousands of subjects, have become increasingly popular in recent years. There have also been efforts to improve the performance of these parameter estimation methods to suit largescale fMRI data [17], [18]. However, in many cases, the neuroimaging data is either located in different places, requiring expensive and time-consuming centralization or is not readily sharable due to privacy or regulatory concerns. In this case, the types of analysis that can be performed are still limited. Many research groups worldwide are collecting comparable data focusing on similar problems. Neuroimaging analysis can be significantly improved when all such worldwide datasets can be combined in more extensive studies.

One approach to perform such larger studies is to combine all the data in a centralized location and perform analysis. If possible, this is a useful approach, however it is inefficient and can present barriers particularly for neuroimaging-based studies for two main reasons. First, MRI data is usually large and saving all the data in one central location not only has high transmission costs and delays, but also requires large amounts of redundant storage. Secondly, in some cases, data cannot be readily shared (which represents a substantial amount of the existing data) due to privacy or regulatory restrictions.

The Enhancing NeuroImaging Genetics through Meta-Analysis (ENIGMA) project addresses this by building large-scale consortia who agree to use shared scripts to run local analyses then share results centrally for a meta analysis[19]. This approach has been widely adopted and is highly successful, but precludes high dimensional studies such as voxel based imaging, or iterative approaches. While some ENIGMA project do indeed centralize raw data for certain projects, in many cases this is not possible. Other approaches to fitting linear mixed models (LMEs) for large-scale neuroimaging data have been proposed. Authors of [20] proposed a method of moments approach to estimating variance components with a linear regression; while this approach allows for flexible random effects models it won’t be as scalable as the maximum likelihood approach employed in big linear mixed models (BLMM) [18]. Recently Fan et al [21] proposed a refined version of this approach, by discretizing the estimated variance components into bins, allowing groups of voxels to be whitened with a common estimated covariance. While this is yet more efficient, it further sacrifices accuracy for speed with with the discretization.

One powerful way to bypass these problems is to use decentralized (or federated) algorithms, which do not require assembling the data in one central location. Decentralized learning algorithms have become popular in recent years due to the demand for collaborative studies involving diverse worldwide datasets without worrying about data transmission or violating privacy. The development of decentralized algorithms is an active research area which has applications in many different domains. In this work, we develop and apply a decentralized LME model and show that decentralized prediction models achieve performance similar to that of its equivalient centralized model. We perform detailed experiments on voxel-based morphometry (VBM) features, namely voxelwise gray matter volume, extracted from structural MRI data and demonstrate the robustness of our approach. We implement our approach within the Collaborative Informatics and Neuroimaging Suite Toolkit for Anonymous Computation (COINSTAC) [22, 23] framework employing Big Linear Mixed Models(BLMM) [18] for our decentralized analysis.

The remainder of this paper is organized as follows. Detailed background on LME, BLMM, COINSTAC and decentralized learning methods are discussed in section 2. Our decentralized LME approach is discussed in section 3, followed by results in section 4, with the conclusion in section 5.

## 2 Preliminaries

In this section, we provide a brief introduction of some of the concepts behind our decentralized LME method.

### 2.1 LME in neuroimaging

LME modeling, an extension of linear regression modeling, is used to describe the relationship between a response variable and independent variables, whilst accounting for grouping factors and covariance structure present in the data. This model consists of fixed and random effects. Fixed effects are the covariates that are assumed to have a true value in the population; random effects are assumed to be generated from a stochastic process, being randomly different for each unit. The most common motivation for mixed effects models is with longitudinal data. Subjects measured repeatedly over time will generate data that is correlated within subject, and naively analyzing with fixed effects models will typically produce inflated false positives[24]. In a LME model, a random intercept for each subject is fit, accounting for intrasubject correlation. However, when there are no repeated measurements, random effects can be useful, for example if we wish to model variation of site means as random. Another common use of random effects is to allow additional variation due to a covariates. For example, if subjects are measured at several visits we can have fixed effect of time (common to all subjects), but also random (subject-specific) effect of time.

In full generality, a LME can contain an arbitrary number of fixed effects and random effects, with the random effects being grouped by multiple factors. Linear mixed effects regression model takes the form *Y* = *Xβ* + *Zb* + *ε* where *Y* is the dependent variable vector, *X* is fixed effects matrix and *Z* is random effects matrix, *β* is fixed effects parameter vector, *b* is the random effects vector and *ε* is the error vector; the random and error effects are assumed to be mutually independent and normally distributed, *b*∼*N* (0, *D*) and *ϵ* ∼*N* (0, *Iσ*^2^). Here, *D* is the random effects covariance matrix, typically assumed to be block diagonal.

LMEs are fit with iterative optimization methods, alternately updating the fixed parameters (*β*) and random covariance parameters (unique elements of D). For details, please refer to [25] which introduced modern LME in this form, and [26] for a comprehensive understanding.

Structural or functional MRI data is commonly used to study underlying brain mechanisms or functional dynamics. When this data is collected over multiple experiments, it inherently contains multi-level covariance structure. Linear mixed-effects models on such data are used to identify such group level covariates.

### 2.2 BLMM

As sMRI/fMRI data is large-scale multi-dimensional(4D) data, it can be computationally challenging to perform regression modeling in this case. While existing neuroimaging software tools exploit vectorization to make ordinary least squares regression extremely fast, they don’t have fully flexible LME capabilities, and existing (non-imaging) LME tools have no facility to vectorise over high dimensional imaging data. This motivates BLMM [18], a Python-based tool for parameter estimation and statistical inference of mass-univariate LMMs for large-scale analysis. BLMM tool utilizes a Fisher Scoring procedure for analysis and efficient vectorization to fit LMEs for many voxels, and is designed for use on high performance computing clusters to analyze large-scale datasets with multiple crossed random factors.

### 2.3 COINSTAC

COINSTAC [22, 23, 27] is a platform created to enable large-scale studies with diverse datasets using decentralization approaches removing the traditional barriers of data-centric approaches. As it can work on the datasets without requiring them to be pooled, COINSTAC solves the problem of researchers wanting to collaborate but being unable to because of data sharing restrictions[28]. COINSTAC allows large-scale analysis of decentralized data with results on par with those that would have been obtained if the data were centralized. It has inbuilt tools developed which performs several decentralized algorithms such as regression, classification and other analysis mainly focused on neuroimaging data. In one example, Gazula et al. [29] used this application to perform a decentralized regression on sMRI data preprocessed using voxel-based morphometry to analyze the structural changes in the brain as linked to age, body mass index, and smoking in several thousand datasets located in three continents. This study also emphasized the benefits of large-scale neuroimaging analysis. COINSTAC implements a wide and growing range of decentralized neuroimaging pipelines and supports such large-scale analysis of decentralized data with results on par with results from pooled data.

### 2.4 Decentralized/federated approach for training ML models

Decentralized learning provides a way for worldwide research groups to collaborate and build more accurate and generalizable prediction models while providing solutions to address data transfer or their data-sharing policies[30]. Also referred to as federated learning in the literature [31], most of the research in decentralized machine learning algorithms is focused on deep learning models due to their flexibility and high performance in a wide number of domains. There have been some other efforts to apply decentralization approaches for simple machine learning algorithms such as support vector machines (SVMs) [32]. In decentralized algorithms, since data cannot be readily shared, all the participating sites (referred as local sites) start with the same machine learning models using the same initial conditions. After each training iteration, each of these local sites transfer their locally learnt parameters to a global/main site for aggregation. The aggregated parameters are then sent to all the local sites to update their models. This step is essential to make sure that all the local models have the same state after each local training iteration. This training step is repeated for several iterations until the model reaches its desired performance or meets its stopping criteria. In the case of decentralized deep learning models, after each iteration, gradients of the models are sent to the main site for aggregation. Another way of obtaining such aggregated model, known as model averaging, is to transfer the entire local model parameters to the main site instead of gradients. The new model is then sent all the local sites which then train on a new batch of data in the next iteration and repeat the process until the model reaches its desired performance. There are several challenges in designing decentralized algorithms, including heterogeneity across different sites, transmission, maintaining synchronization during training iterations, and preserving data privacy. The authors of [33, 34] provide a detailed summary of the machine learning algorithms, challenges, available architectures for decentralization.

## 3 Decentralized LME

Decentralized learning has recently gained attention as it is privacy-preserving and can handle largescale analysis[30]. As mentioned earlier, in decentralized machine learning algorithms, either the training parameters(gradients) or model parameters are shared after each training iteration so that the actual data is not shared but the information required to build an aggregated model is exchanged. In the case of decentralized brain-age prediction which is similar to regression, a decentralized support vector regression algorithm was employed [32] where the support vector model parameters were exchanged to build an aggregated regression model.

Building decentralized LME models for neuroimaging data is not as straightforward as machine learning algorithms because LME modelling consists of two stages, namely, parameter estimation and statistical inference. In an sMRI or fMRI analysis, parameters must be estimated for every voxel in the analysis mask. As there are typically hundeds of thousands of voxels in an image, and parameter estimation for the LME requires computationally intensive numerical methods, such an approach accrues large overheads in computation time and storage. Similarly, the statistical inference step also has the same challenge for voxel-based data. We employ BLMM [18] which models mass-univariate fMRI using a vectorized LME modeling approach. Since the data cannot be shared outside local sites in a decentralized setting, one needs to identify a way to sharing these mass-univariate model parameters, their statistical inference values and also extract information from the data at local sites.

The LME model, for observation *j* at site *i* can be represented in the following form:

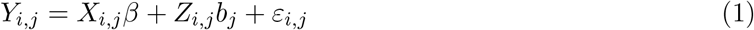

where *X*_*i,j*_ and *Z*_*i,j*_ are (1×*p*) and (1×*q*) row vectors of fixed and random effects covariates for observation *j* at site *i*, respectively. The random terms in the model are *b*_*j*_, the random effects for site *i* and *ε*_*i,j*_, the error term for observation *j* at site *i*. The model parameters are: *β*, the (*p*×1) column vector of fixed effects parameters, *σ*^2^, the scalar variance term, and *D*, the (*q*×*q*) random effects covariance matrix. This model is solved to estimate unknown parameters *β, σ*^2^ and the unique elements of random effects covariance matrix, *D* at each site *i* individually and are then used at the aggregator site to compute the global *β, σ*^2^ and *D* values.

Neuroimaging data analyses are performed directly on the MRI data or on the features extracted from the MRI data. Since these represent different types of data, in this work we propose two decentralized algorithms, one for FreeSurfer features extracted from MRI data and other while working directly with the MRI data. Algorithm-1 outlines all the steps of decentralized LME in both the cases.

### 3.1 MRI Extracted FreeSurfer Features

Our decentralized LME algorithm performs two iterations between local and master sites to obtain an aggregated model which has performance on par with that of a centralized model. FreeSurfer features contain volumes of specific brain structures that can be represented as features.

To perform LME, we form *X*_*i*_, *Y*_*i*_, *Z*_*i*_ matrices at each site *i* combining all observations *j* at that each site to represent *X*_*i,j*_, *Y*_*i,j*_, *Z*_*i,j*_, defined in eq. (1), respectively. Here, *X*_*i*_ and *Z*_*i*_ are (*n*×*p*) and (*n*×*q*) matrices of fixed and random effects covariates of all the observations at each site *i*.

In the first iteration (Iteration-1 in Fig. 1), each local site loads the features of all the subjects present at that site and form *X*_*i*_, *Y*_*i*_, *Z*_*i*_ matrices at each site *i* which represent fixed covariates, FreeSurfer feature data and random effects design matrix respectively. We then employ the BLMM tool to generate parameter estimates and calculate inference for the above matrices. The number of levels of the random factor, the number of local samples(or subjects), the computed *β*_*i*_ and other parameters are sent to the master site from each local site. The master site receives this information from each local site, aggregates them and sends the total number of levels and total observations to all the local sites.

**Figure 1:**
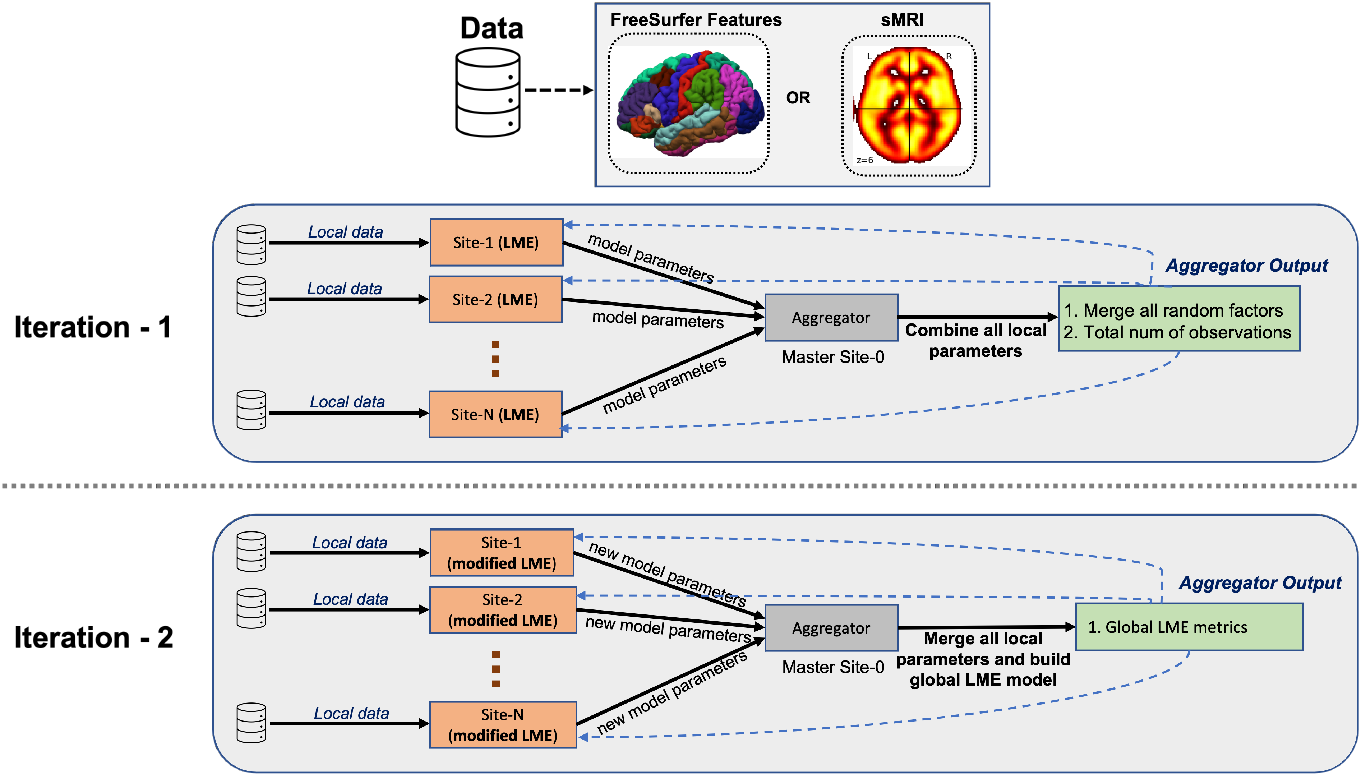
Overall flow of data measures and model parameters from locally trained LME models sent to the main site for aggregation. The aggregated model output from each iteration (marked in dashed blue lines) is sent to all the local sites. The models use the output from the aggregator, transform the data, build local LME models, and send these parameters to the aggregator site. In the final step, the aggregator builds the global LME model and sends model stats to all the local sites.

In the second iteration (Iteration-2 in Fig. 1), each local site recomputes the random effects design matrix, *Z*_*i*_, based on the total number of levels and total observations received from the master site. Using the new *Z*_*i*_, the product matrices *X*_*i*_^*⊤*^*X*_*i*_, *X*_*i*_^*⊤*^*Y*_*i*_, *X*_*i*_^*⊤*^*Z*_*i*_, *Y*_*i*_^*⊤*^*X*_*i*_, *Y*_*i*_^*⊤*^*Y*_*i*_, *Y*_*i*_^*⊤*^*Z*_*i*_, *Z*_*i*_^*⊤*^*X*_*i*_, *Z*_*i*_^*⊤*^*Y*_*i*_, *Z*_*i*_^*⊤*^*Z*_*i*_ are computed at each site *i* and are sent to the master site. These product matrices transform the actual data to another dimension space and therefore serve as an encrypted data and can be sent to other sites. Also, as these product matrices have lower dimensions than the original data, they can be transmitted without heavy bandwidth requirement. At the master site, global product matrices are generated by adding the corresponding matrix received from all the local sites as shown below.

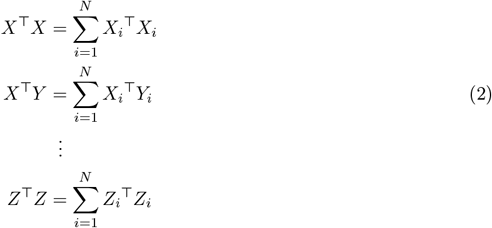

The master site then generates parameter estimates and calculates statistical inference values using the above global product matrices. All the global estimates along with the inference values are sent to the local sites. The master site has the aggregated LME model constructed using a decentralized approach.

### 3.2 MRI data

Our decentralized LME algorithm performs three iterations between local and master sites to obtain an aggregated model. For simplicity, let us assume that all the sMRI or fMRI data is represented in the `nifti (.nii)` format [35], [36].

In the initial iteration (Iteration-0 in Fig. 2), each local site loads the `nifti` data of all the subjects present at that site, computes average of the `nifti` images and shares this subject-wise average `nifti` image to the master site. At the master site, it gathers all these subject-wise average `nifti` files, computes a mean image of all the average nifti files received from all the local sites and filters all the voxel values using a threshold to get a mean mask image. It then constructs a new mask image from mean mask image using MNI template[37], [38], which serves as a mask and is sent to all the local sites. Computing such a common mask is important to make sure that data across all the sites is filtered using the same mask file and the mask file need to be in compliance with the nifti data from all the sites.

**Figure 2:**
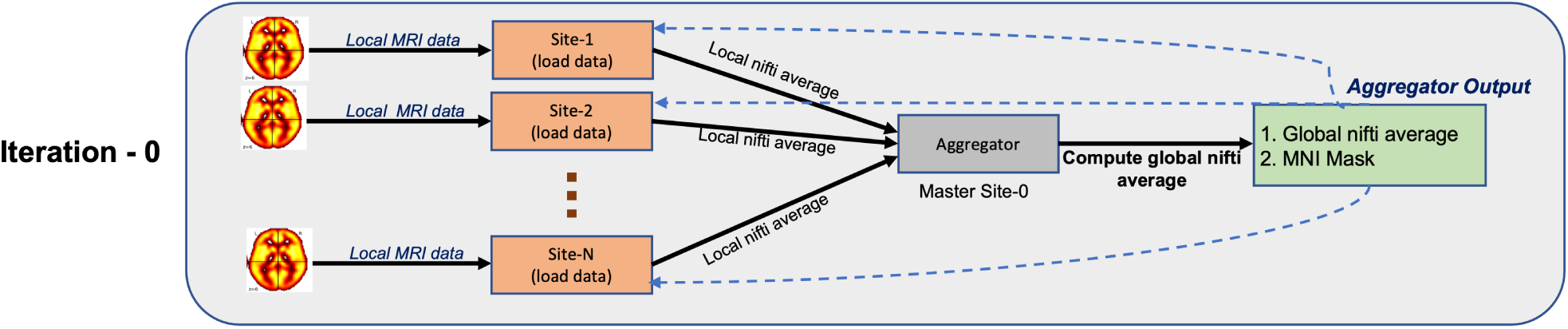
Initial iteration performed on MRI data to generate a uniform mask image to be used across all the sites to load `nifti` files.

The next two iterations follow same steps as that of above decentralized FreeSurfer feature LME model. In the next iteration (Iteration-1 in Fig. 1), all the local sites reload the data using the common new mask sent by the master site and form *X*_*i*_, *Y*_*i*_, *Z*_*i*_ matrices at each site *i* which represent fixed covariates, nifti data and random effects design matrix respectively. To improve the computation performance of estimation, we compute product of *X*_*i*_, *Y*_*i*_, *Z*_*i*_ and their transposes *X*_*i*_^*⊤*^, *Y*_*i*_^*⊤*^, *Z*_*i*_^*⊤*^ to generate *X*_*i*_^*⊤*^*X*_*i*_, *X*_*i*_^*⊤*^*Y*_*i*_, *X*_*i*_^*⊤*^*Z*_*i*_, *Y*_*i*_^*⊤*^*X*_*i*_, *Y*_*i*_^*⊤*^*Y*_*i*_, *Y*_*i*_^*⊤*^*Z*_*i*_, *Z*_*i*_^*⊤*^*X*_*i*_, *Z*_*i*_^*⊤*^*Y*_*i*_, *Z*_*i*_^*⊤*^*Z*_*i*_. These product matrices have lower dimensions compared to the original matrices and therefore have lower memory consumption. We then employ BLMM tool to generate parameter estimates and calculate inference for the above product matrices. The number of levels of random factor, the number of local samples(or subjects), the computed *β*_*i*_ and other parameters are sent to the master site from each local site. The master site receives this information from each local site, aggregates them and sends the total number of levels and total observations to all the local sites.

In the final iteration (Iteration-2 in Fig. 1), each local site generates the random effects design matrix, *Z*_*i*_, based on the total number of levels and total observations received from the master site. Using the new *Z*_*i*_, the product matrices *X*_*i*_^*⊤*^*X*_*i*_, *X*_*i*_^*⊤*^*Y*_*i*_, *X*_*i*_^*⊤*^*Z*_*i*_, *Y*_*i*_^*⊤*^*X*_*i*_, *Y*_*i*_^*⊤*^*Y*_*i*_, *Y*_*i*_^*⊤*^*Z*_*i*_, *Z*_*i*_^*⊤*^*X*_*i*_, *Z*_*i*_^*⊤*^*Y*_*i*_, *Z*_*i*_^*⊤*^*Z*_*i*_ are recomputed at each site *i* and these product matrices are sent to the master site. The product matrices transform the actual data to another dimension space and therefore serves as an encrypted data and can be sent to other sites. Also, as these product matrices have lower dimensions than the original data and do not scale with *n*, the number of observations,, they can be transmitted without heavy bandwidth requirement. At the master site, global product matrices are generated by adding the corresponding matrix received from all the local sites as shown in Eq.(2). The aggregator site computes the global parameter estimates and calculates the statistical inference values using the global product matrices. All the global parameter estimates along with the statistical results are sent to the local sites. The master site has now aggregated LME model using a decentralized approach.

### 3.3 Design specifications for inputs

As there is a lot of heterogeneity involved in decentralized computations, below are some of the design specifications for our input matrices across all the sites.

X is fixed effects matrix where the number of columns corresponds to the number of fixed effects and number of rows correspond to number of observations. The number of fixed effects remain the same across all the participating local sites whereas number of rows can be different across sites.

Y is vector of dependent variables which can correspond to any neuroimaging features. If this corresponds to FreeSurfer stats, the number of such stat variables should be the same across all the local sites though the number of subjects can vary. If it is VBM voxel data, the voxel dimension of the nifti file of all the subjects and across the sites should be same.

*Z* is the random effects covariates matrix. In our design, we consider only one random factor, that is site covariate, which is a categorical variable indicating the site it is from. So the number of random effects for site factor is just 1. On deciding the number of levels, each local site has sub-sites which forms the site covariate for that particular local site. Suppose the local site 1 has 5 sub-sites, then the Z matrix for that site has to be framed accordingly based on the sub-site data. There are two cases which handle sub-site data across multiple local sites, either the sub-sites across local sites are entirely different or some of the local sites share same sub-sites. In the first case when the sub-sites across the local sites are entirely different, lets suppose the local site 1 has 3 sub-sites *l*_1_, *l*_2_, *l*_3_ and the local site 2 has 3 sub-sites which are different from sub-sites of the local site 1, say *l*_4_, *l*_5_, *l*_6_ then the *Z* matrix formed in the third iteration of the algorithm(1) has 6 different columns as we stack the *Z*_*i*_ matrices from all local sites to solve LME in the central node in COINSTAC. In the other case of dealing with shared sub-sites across local sites, suppose local site 1 has sub-sites *l*_1_, *l*_2_, *l*_3_ and local site 2 has sub-sites *l*_1_, *l*_2_ and *l*_4_, then the *Z* matrix has to be formed differently with 4 columns which would require all the sites has to follow a standardized sub-site names. In the current implementation, we assume that all the local sites have data from different sub-sites.

#### Implementation in COINSTAC

COINSTAC provides an open platform to implement new decentralized algorithms. It has inbuilt tools developed which can be used to perform several decentralized algorithms such as regression, classification and other analysis mainly focused on neuroimaging data. Fig. 3 demonstrates the execution of decentralized LME model. In this implementation, the main functionalities of local sites involve forming *X*_*i*_,*Y*_*i*_ and *Z*_*i*_ matrices from the inputs, calculate corresponding product matrices, estimate parameter estimates and inference results using PSFS method. The generation of *Z*_*i*_ matrix can be complicated as it involves assumptions regarding random effects and levels. The number of random effects per factor should be agreed upon and decided beforehand, as this is a basic design question. In our current implementation, the number of levels can vary but all the local sites should be using the same format for *Z*_*i*_. Based on the provided inputs, each local site generates the global random effects design matrix(*Z*) including random factors and effects from all local sites(in the iteration-2 of the algorithm 1). The aggregator site mainly calculates the total number of levels and total observations consolidated from all local sites, performs parameter estimation for decentralised LME analysis using the PSFS algorithm and outputs contrast images from all local sites to calculate statistical inference results.

**Figure 3:**
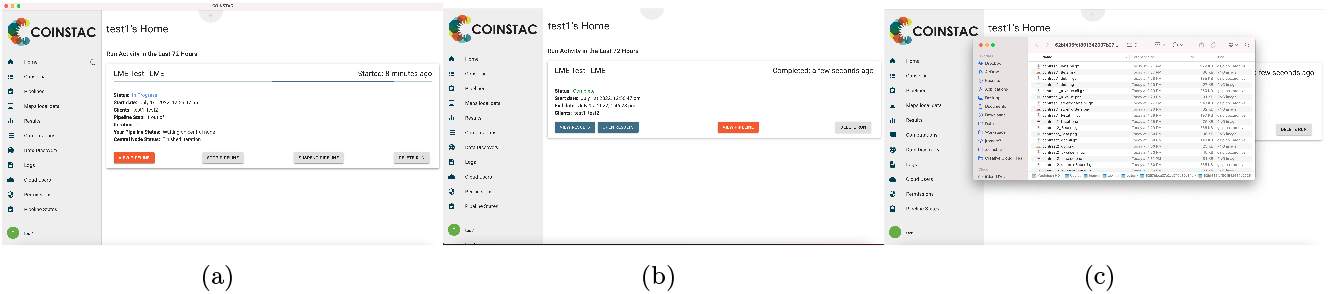
Screenshots of the COINSTAC tool running decentralized LME model. (a) shows the screen where decentralized LME is running in COINSTAC GUI and (b) shows the completion screen with the results files displayed in (c).

##### Algorithm 1: Decentralized LME Regression

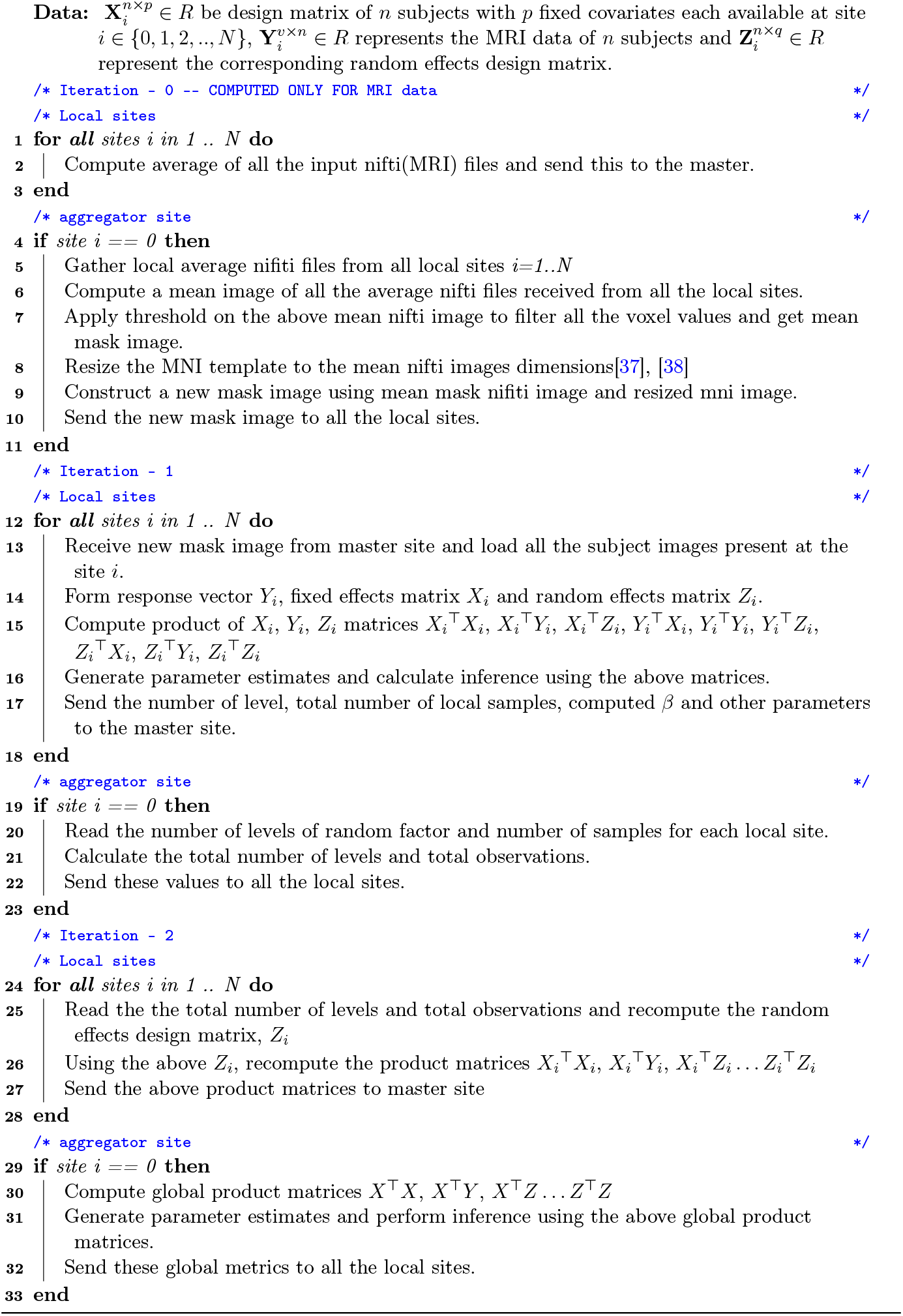

### 3.4 Metrics

We use the following metrics to determine the effectiveness of our approach. We compute these metrics for centralized model where all the data is pooled in one location. These metrics are also for the aggregated LME model trained in a decentralized fashion. We then compare metric values for both models.

**Residual mean squares** is one of the most commonly used metrics in regression models to compute the residual error. The lower the value of this metric, the better a model fits a given dataset. Residual mean squares for a *β* estimate is defined as follows:

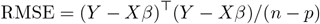

where *β* is an estimate of the parameter vector, *n* is the number of observations/input niftis (potentially spatially varying) and *p* is the number of fixed effects parameters.

The **log-likelihood** value of a regression model provides a way to measure how good a model fits a dataset. The higher the value of this metric, the better a model fits a dataset. The log likelihood of *β, σ*^2^, *D* is computed as follows:

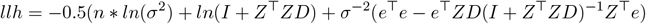

where *n* is the total number of observations (potentially spatially varying), *Z* is the random effects covariates matrix, *e* is the ordinary least square(OLS) residuals (*e* = *Y − Xβ*), *σ*^2^ is the fixed effects variance and *D* is the random effects covariance matrix.

## 4 Experiments & Results

In this section, we implement the decentralized LME in COINSTAC and solve the LME regression model of the form *Y*_*i*_ = *X*_*i*_*β* + *Z*_*i*_*b* + *ε*_*i*_ using Pseudo Simplified Fisher Scoring (PSFS) method in BLMM tool [39]. All the decentralized computations discussed here have been performed on a single machine.

### 4.1 Data

T1-weighted *s*MRI images of 98 chronic schizophrenic patients and 99 controls, taken from the MIND Clinical Imaging Consortium (MCIC), were used for this experiment. For additional information about the data and preprocessing, please refer to [40]. In our LME analysis, age, diagnosis, and gender are used as fixed covariates of the model, and the site covariates are used as random effects. We test our decentralized LME model with two sites with one site having 52 Controls/52 patients and the other 47 controls/46 patients respectively.

### 4.2 FreeSurfer features

FreeSurfer (version v5.3) [41] was used to extract brain structural features from the *s*MRI data. A standard `aseg.stats` file that has features corresponding to total intracranial volume (eTIV), left hemisphere (lh) and right hemisphere (rh) subcortical regions is generated. Volume features corresponding to 7 regions are used as FreeSurfer features for analysis. For a site *i*, the FreeSurfer features serve as the *Y*_*i*_ matrix, the fixed covariates form the *X*_*i*_ matrix, and the site covariates are used to design random effects *Z*_*i*_ matrix.

We compare decentralized and centralized values for *res*_*ms* and *llh* metrics as shown in Fig. 4. It can be seen that decentralized values are similar to centralized values for both the metrics. Additionally, we plot correlations of LME model parameters (*β* values of the fixed covariates) of centralized LME model on X-axis and of decentralized LME model on Y-axis and compare it on an identity line (*Y* = *X*). In Fig.7, we plot a reference identity line in green and plot these beta coordinates in red. As shown in the figure 5, most of the points follow the reference line showing that our decentralized model has parameters similar to the centralized model.

**Figure 4:**
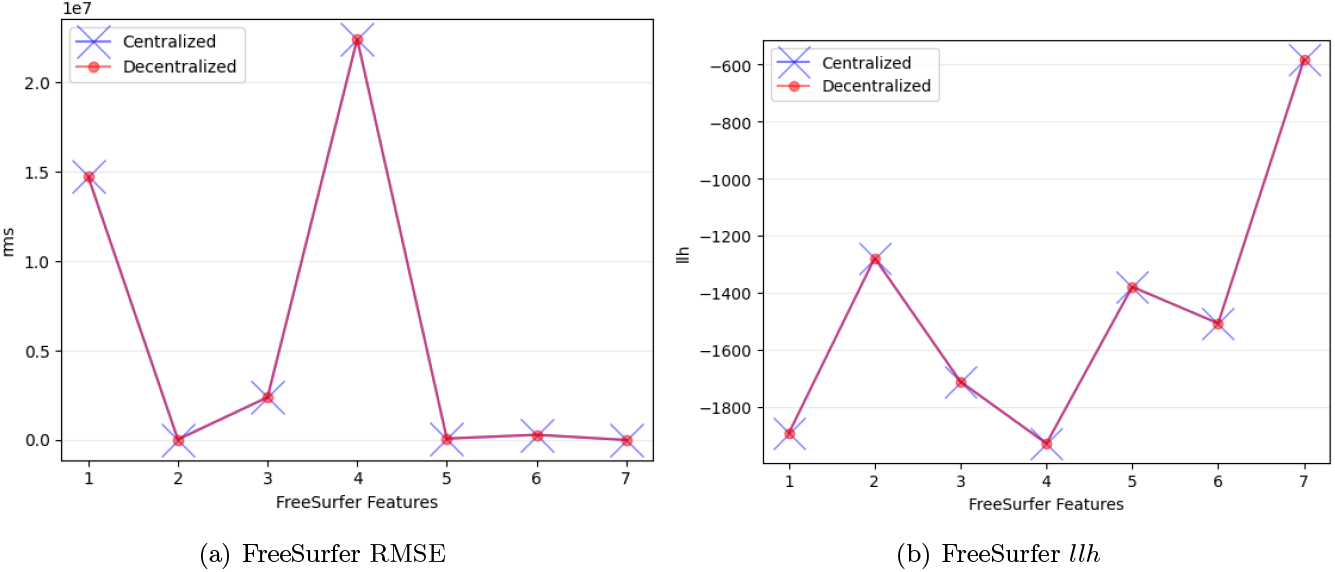
Comparison of decentralized and centralized values for RMSE and *llh* metrics.

**Figure 5:**
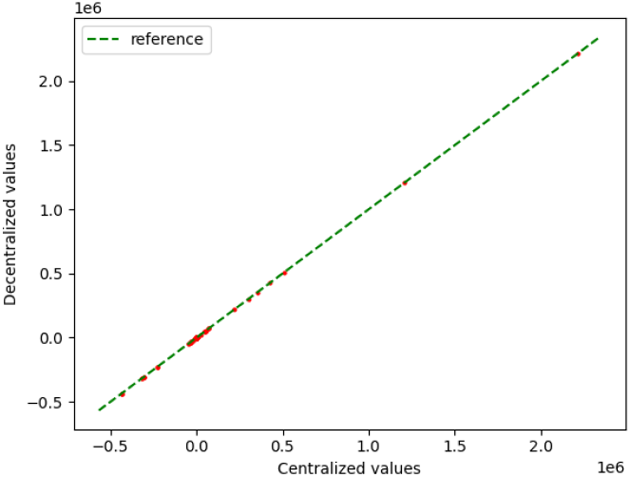
Scatterplot of *β* values with centralized on X-axis and decentralized on Y-axis.

### 4.3 MRI data

The dimensions of the VBM stats are the same across all the sites for all observations. For a site *i*, the resulting in-brain MNI data serves as the *Y*_*i*_ matrix, the fixed covariates form the *X*_*i*_ matrix, and the site covariates are used to design random effects *Z*_*i*_ matrix.

To validate the LME models, we plot voxel-wise *β* values corresponding to the *isControl* covariate of both centralized (fig. 6(a)) and decentralized (fig. 6(b)) LME models. In Fig. 6, we observe that controls show larger differences in bilateral insula, temporal lobe, medial frontal lobe, and cerebellum, as expected.

**Figure 6:**
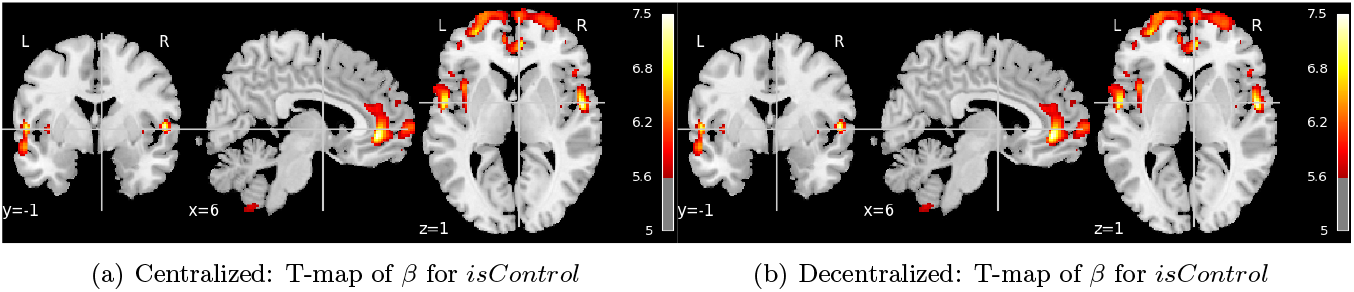
Rendered image showing T-map of *β* values corresponding to the *isControl* covariate.

Furthermore, we plot (fig.7) the scatterplot of *β*’s corresponding to fixed covariates (age, isControl, gender) with the X-axis representing the centralized model and Y corresponding to the decentralized model. We also plot a reference identity line (correlation = 1). As shown in the figure (fig. 7), most of the points follow the reference line showing that our decentralized model has fitted parameters similar to the centralized model. Fig. 8 and fig. 9 are the nifti images corresponding to residual mean squares(*res*_*ms*) and log-likelihood (*llh*) metrics respectively. The scatterplot of these metrics shown in fig. 8(c) and fig. 9(c) also have similar behavior around 0.

**Figure 7:**
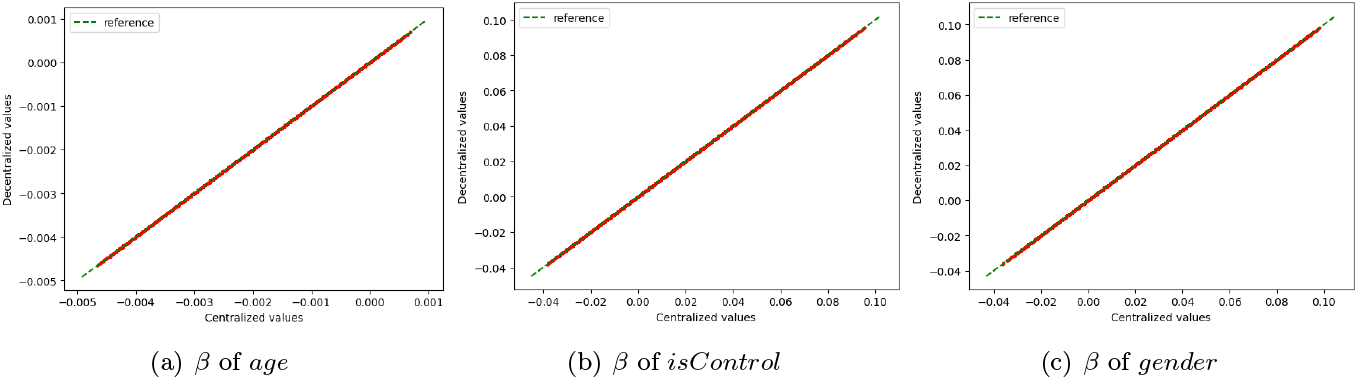
Comparison of *β* values corresponding to fixed covariates of the model. For each voxel, the centralized *β* value is considered as X-coordinate and the corresponding decentralized *β* is considered as Y-coordinate.

**Figure 8:**
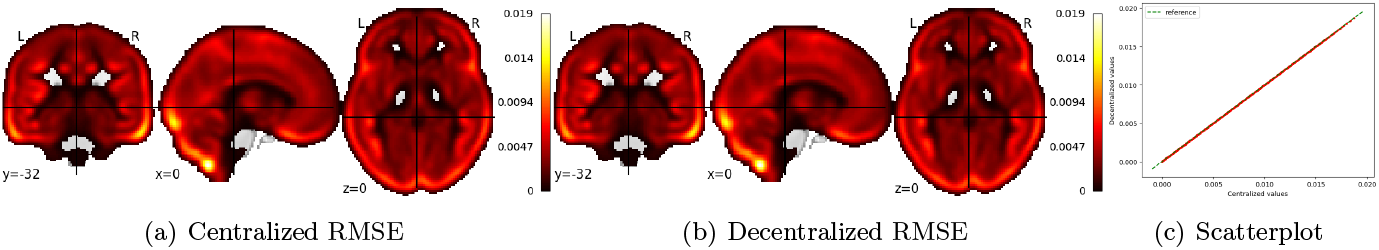
Snapshot of a nifti image showing the residual mean squares (a), (b) and scatterplot of these values with centralized on X-axis and decentralized metrics on Y-axis.

**Figure 9:**
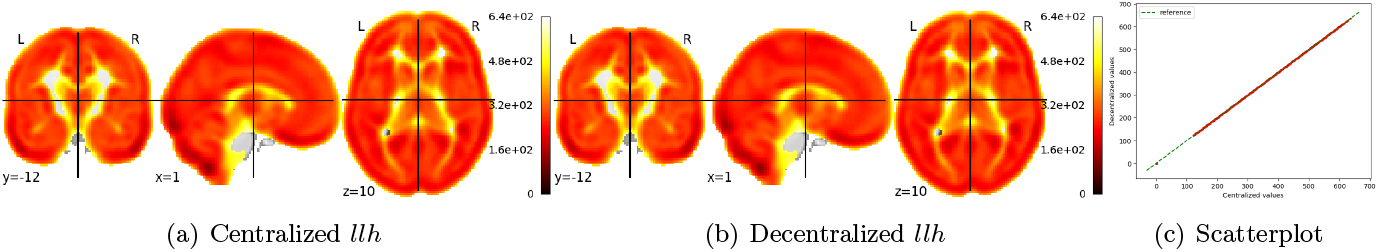
Snapshot of a nifti image showing the log-likelihood metrics.

To statistically compare the performance of centralized and decentralized models for these *β* values, we performed a *t*-test with a null hypothesis that the differences between two samples have a distribution centered about zero. Results (with *α* = 0.05) fail to reject the null hypothesis, thus suggesting that the decentralized models achieve performance similar to that of centralized models.

## 5 Conclusion

In this work, we propose a decentralized approach to model LME regression and compare the results with a centralized LME model. The decentralized model is built utilizing the information from local LME models at different sites and involves no data sharing. Results from models trained using sMRI VBM data show that the performance of the decentralized model is as good as the centralized model. Decentralization provides us a way to perform larger studies without the need to share the actual data encouraging different research groups to readily collaborate in a way that is compliant with data-sharing policies and minimizes data transmission and computation resources.

## 6 Acknowledgements

This work was funded by NIH R01DA040487 and NSF 2112455.

## Notes

### Competing Interest Statement

The authors have declared no competing interest.

## References

[1] C. Beckmann, M. Jenkinson, and S. Smith, “General multilevel linear modeling for group analysis in fmri,” NeuroImage, vol. 20, p. 1052–63, 10 2003.

[2] M. W. Woolrich, T. E. Behrens, C. F. Beckmann, M. Jenkinson, and S. M. Smith, “Multilevel linear modelling for fmri group analysis using bayesian inference,” NeuroImage, vol. 21, no. 4, p. 1732 –1747, 2004.

[3] K. Friston, K. Stephan, T. Lund, A. Morcom, and S. Kiebel, “Mixed-effects and fmri studies,” NeuroImage, vol. 24, p. 244–52, 02 2005.

[4] J. L. Bernal-Rusiel, D. N. Greve, M. Reuter, B. Fischl, and M. R. Sabuncu, “Statistical analysis of longitudinal neuroimage data with linear mixed effects models,” NeuroImage, vol. 66, p. 249–260, 2013.

[5] G. Chen, Z. Saad, J. Britton, D. Pine, and R. Cox, “Linear mixed-effects modeling approach to fmri group analysis.,” NeuroImage, vol. 73, 01 2013.

[6] T. Madhyastha, M. Peverill, N. Koh, C. McCabe, J. Flournoy, K. Mills, K. King, J. Pfeifer, and K. A. McLaughlin, “Current methods and limitations for longitudinal fmri analysis across development,” Developmental Cognitive Neuroscience, vol. 33, p. 118 –128, 2018. Methodological Challenges in Developmental Neuroimaging: Contemporary Approaches and Solutions.

[7] T. K. Koerner and Y. Zhang, “Application of linear mixed-effects models in human neuroscience research: a comparison with pearson correlation in two auditory electrophysiology studies,” Brain sciences, vol. 7, no. 3, p. 26, 2017.

[8] Z. Yu, M. Guindani, S. F. Grieco, L. Chen, T. C. Holmes, and X. Xu, “Beyond t test and anova: applications of mixed-effects models for more rigorous statistical analysis in neuroscience research,” Neuron, 2021.

[9] N. Lange, “What can modern statistics offer imaging neuroscience?,” Statistical methods in medical research, vol. 12, no. 5, p. 447–469, 2003.

[10] G. Chen, Z. S. Saad, J. C. Britton, D. S. Pine, and R. W. Cox, “Linear mixed-effects modeling approach to fmri group analysis,” Neuroimage, vol. 73, p. 176–190, 2013.

[11] J. A. Mumford and T. Nichols, “Modeling and inference of multisubject fmri data,” IEEE Engineering in Medicine and Biology Magazine, vol. 25, no. 2, p. 42–51, 2006.

[12] C. F. Beckmann, M. Jenkinson, and S. M. Smith, “General multilevel linear modeling for group analysis in fmri,” Neuroimage, vol. 20, no. 2, p. 1052–1063, 2003.

[13] J. A. Mumford and R. A. Poldrack, “Modeling group fmri data,” Social cognitive and affective neuroscience, vol. 2, no. 3, p. 251–257, 2007.

[14] D. Szucs and J. P. Ioannidis, “Sample size evolution in neuroimaging research: An evaluation of highly-cited studies (1990–2012) and of latest practices (2017–2018) in high-impact journals,” NeuroImage, vol. 221, p. 117164, 2020.

[15] B. J. Casey, T. Cannonier, M. I. Conley, A. O. Cohen, D. M. Barch, M. M. Heitzeg, M. E. Soules, T. Teslovich, D. V. Dellarco, H. Garavan, et al., “The adolescent brain cognitive development (abcd) study: imaging acquisition across 21 sites,” Developmental cognitive neuroscience, vol. 32, p. 43–54, 2018.

[16] C. Sudlow, J. Gallacher, N. Allen, V. Beral, P. Burton, J. Danesh, P. Downey, P. Elliott, J. Green, M. Landray, et al., “Uk biobank: an open access resource for identifying the causes of a wide range of complex diseases of middle and old age,” PLoS medicine, vol. 12, no. 3, p. e1001779, 2015.

[17] K. J. Friston, K. E. Stephan, T. E. Lund, A. Morcom, and S. Kiebel, “Mixed-effects and fmri studies,” Neuroimage, vol. 24, no. 1, p. 244–252, 2005.

[18] T. Maullin-Sapey and T. Nichols, “Blmm: Parallelised computing for big linear mixed models,” bioRxiv, 2022.

[19] C. E. Bearden and P. M. Thompson, “Emerging global initiatives in neurogenetics: the enhancing neuroimaging genetics through meta-analysis (enigma) consortium,” Neuron, vol. 94, no. 2, p. 232–236, 2017.

[20] M. A. Lindquist, J. Spicer, I. Asllani, and T. D. Wager, “Estimating and testing variance components in a multi-level glm,” NeuroImage, vol. 59, no. 1, p. 490–501, 2012.

[21] C. C. Fan, C. E. Palmer, J. R. Iversen, D. Pecheva, D. Holland, O. Frei, W. K. Thompson, D. J. Hagler Jr, O. A. Andreassen, T. L. Jernigan, et al., “Fema: Fast and efficient mixed-effects algorithm for population-scale whole-brain imaging data,” BioRxiv, p. 2021–10, 2021.

[22] S. M. Plis, A. D. Sarwate, D. Wood, C. Dieringer, D. Landis, C. Reed, S. R. Panta, J. A. Turner, J. M. Shoemaker, K. W. Carter, et al., “Coinstac: a privacy enabled model and prototype for leveraging and processing decentralized brain imaging data,” Frontiers in neuroscience, vol. 10, p. 365, 2016.

[23] “COINSTAC.” http://coinstac.trendscenter.org.

[24] J. A. Mumford and R. A. Poldrack, “Modeling group fMRI data,” Social Cognitive and Affective Neuroscience, vol. 2, p. 251–257, 09 2007.

[25] N. M. Laird and J. H. Ware, “Random-effects models for longitudinal data,” Biometrics, p. 963–974, 1982.

[26] J. Pinheiro and D. Bates, Mixed-effects models in S and S-PLUS. Springer science & business media, 2006.

[27] T. White, E. Blok, and V. D. Calhoun, “Data sharing and privacy issues in neuroimaging research: Opportunities, obstacles, challenges, and monsters under the bed,” Human Brain Mapping, 2020.

[28] J. Ming, E. Verner, A. Sarwate, R. Kelly, C. Reed, T. Kahleck, R. Silva, S. Panta, J. Turner, S. Plis, et al., “Coinstac: Decentralizing the future of brain imaging analysis,” F1000Research, vol. 6, 2017.

[29] H. Gazula, B. Holla, Z. Zhang, J. Xu, E. Verner, R. Kelly, G. Schumann, and V. D. Calhoun, “Decentralized multi-site vbm analysis during adolescence shows structural changes linked to age, body mass index, and smoking: A coinstac analysis,” bioRxiv, p. 846386, 2019.

[30] A. D. Sarwate, S. M. Plis, J. A. Turner, M. R. Arbabshirani, and V. D. Calhoun, “Sharing privacy-sensitive access to neuroimaging and genetics data: a review and preliminary validation,” Frontiers in neuroinformatics, vol. 8, p. 35, 2014.

[31] K. Rootes-Murdy, H. Gazula, E. Verner, R. Kelly, T. DeRamus, S. Plis, A. Sarwate, J. Turner, and V. Calhoun, “Federated analysis of neuroimaging data: A review of the field,” Neuroinformatics, vol. 20, no. 2, p. 377–390, 2022.

[32] S. Basodi, R. Raja, B. Ray, H. Gazula, A. D. Sarwate, S. Plis, J. Liu, E. Verner, and V. D. Calhoun, “Decentralized brain age estimation using mri data,” Neuroinformatics, p. 1–10, 2022.

[33] M. Aledhari, R. Razzak, R. M. Parizi, and F. Saeed, “Federated learning: A survey on enabling technologies, protocols, and applications,” IEEE Access, vol. 8, p. 140699–140725, 2020.

[34] T. Li, A. K. Sahu, A. Talwalkar, and V. Smith, “Federated learning: Challenges, methods, and future directions,” IEEE Signal Processing Magazine, vol. 37, no. 3, p. 50–60, 2020.

[35] M. Larobina and L. Murino, “Medical image file formats,” Journal of digital imaging, vol. 27, no. 2, p. 200–206, 2014.

[36] X. Li, P. S. Morgan, J. Ashburner, J. Smith, and C. Rorden, “The first step for neuroimaging data analysis: Dicom to nifti conversion,” Journal of neuroscience methods, vol. 264, p. 47–56, 2016.

[37] V. S. Fonov, A. C. Evans, R. C. McKinstry, C. Almli, and D. Collins, “Unbiased nonlinear average age-appropriate brain templates from birth to adulthood,” NeuroImage, vol. 47, p. S102, 2009.

[38] V. Fonov, A. C. Evans, K. Botteron, C. R. Almli, R. C. McKinstry, D. L. Collins, B. D. C. Group, et al., “Unbiased average age-appropriate atlases for pediatric studies,” NeuroImage, vol. 54, no. 1, p. 313–327, 2011.

[39] T. Maullin-Sapey and T. E. Nichols, “Fisher scoring for crossed factor linear mixed models,” Statistics and computing, vol. 31, no. 5, p. 1–25, 2021.

[40] R. L. Gollub, J. M. Shoemaker, M. D. King, T. White, S. Ehrlich, S. R. Sponheim, V. P. Clark, J. A. Turner, B. A. Mueller, V. Magnotta, et al., “The mcic collection: a shared repository of multi-modal, multi-site brain image data from a clinical investigation of schizophrenia,” Neuroinformatics, vol. 11, no. 3, p. 367–388, 2013.

[41] B. Fischl, “Freesurfer,” Neuroimage, vol. 62, no. 2, p. 774–781, 2012.

